## Supplementary Information for "Single trial dynamics of attentional intensity in visual area V4"

### Supplementary Table

**Supplementary Table S1. Modulation indices of attentional intensity.** Modulation indices of attentional intensity on behavioral and neural correlates, computed across intensity conditions.

| **Modulation Index** | **Mean ± SEM** | **p-value** |
|---|---|---|
| Reward (N = 24) | 0.598 ± 0.015 | $p < 10^{-21}$ |
| Pupil area (N = 24) | 0.102 ± 0.013 | $p < 10^{-7}$ |
| Behavioral d’ (N = 24) | 0.265 ± 0.011 | $p < 10^{-16}$ |
| Microsaccades towards stimulus location (0-200 ms, N = 24) | –0.046 ± 0.068 | p = 0.53 |
| Neuronal d’ (n = 970) | 0.130 ± 0.004 | $p < 10^{-153}$ |
| Fano factor | –0.009 ± 0.001 | $p < 10^{-26}$ |
| Spike-count correlation | –0.037 ± 0.003 | $p < 10^{-30}$ |

**Supplementary Table S2: Mean behavioral hit and false alarm (FA)**

| | Hit rate (%) (Mean ± SEM) | | False alarm rate (%) (Mean ± SEM) | |
|---|---|---|---|---|
| | Small reward | Large reward | Small reward | Large reward |
| Monkey P (N = 9) | 85.9 ± 2.2 | 92.8 ± 0.7 | 34.7 ± 2.4 | 13.1 ± 1.4 |
| Monkey S (N = 15) | 79.1 ± 1.4 | 89.1 ± 1.0 | 30.3 ± 1.4 | 15.4 ± 1.3 |

**Supplementary Table S3: Spatial RF location and size**

| | RF eccentricity (°) Mean ± SEM | RF polar angle(°) Mean ± SEM | RF size, sigma (°) Mean ± SEM |
|---|---|---|---|
| Monkey P (n = 66) | 2.8 ± 0.1 | 132.4 ± 2.6 | 1.13 ± 0.05 |

| | | | |
|---|---|---|---|
| Monkey S (n = 81) | 2.7 ± 0.1 | 108.8 ± 3.2 | 1.18 ± 0.06 |

**Supplementary Table S4: Modulation of behavioral, physiological and neurophysiological responses during early and late trials within a session.**

| | Monkey P | | | | | |
|---|---|---|---|---|---|---|
| | Reward size | Early trials | Late trials | ANOVA | | |
| | | | | Attention intensity | Trial timing | Intensity-by-time interaction |
| Behavioral d’ | Small | 1.51 ± 0.13 | 1.59 ± 0.11 | $F(1,32) = 100.94$ ($p < 10^{-4}$) | $F(1,32) = 2.96$ ($p = 0.09$) | $F(1,32) = 0.7$ ($p = 0.4$) |
| | Large | 2.45 ± 0.08 | 2.72 ± 0.08 | | | |
| %Abort trials | Small | 46.9 ± 3.5 | 56.3 ± 4.2 | $F(1,32) = 36.9$ ($p < 10^{-4}$) | $F(1,32) = 3.41$ ($p = 0.07$) | $F(1,32) = 1.43$ ($p = 0.24$) |
| | Large | 31.9 ± 1.9 | 33.9 ± 1.9 | | | |
| Pupil area | Small | 1.043 ± 0.007 | 0.947 ± 0.011 | $F(1,159) = 108.86$ ($p < 10^{-4}$) | $F(1,159) = 160.26$ ($p < 10^{-4}$) | $F(1,159) = 1.77$ ($p = 0.18$) |
| | Large | 1.143 ± 0.014 | 1.027 ± 0.008 | | | |
| V4 spike rate | Small | 0.807 ± 0.008 | 0.764 ± 0.010 | $F(1,16267) = 67.84$ ($p < 10^{-15}$) | $F(1,16267) = 67.44$ ($p < 10^{-15}$) | $F(1,16267) = 3.01$ ($p = 0.08$) |
| | Large | 0.927 ± 0.006 | 0.843 ± 0.009 | | | |
| | Monkey S | | | | | |
| | Reward size | Early trials | Late trials | ANOVA | | |
| | | | | Attention intensity | Trial timing | Intensity-by-time interaction |
| Behavioral d’ | Small | 1.47 ± 0.07 | 1.25 ± 0.07 | $F(1,56) = 162.6$ ($p < 10^{-4}$) | $F(1,56) = 1.38$ ($p = 0.24$) | $F(1,56) = 3.55$ ($p = 0.06$) |
| | Large | 2.26 ± 0.07 | 2.31 ± 0.07 | | | |
| %Abort trials | Small | 54.4 ± 2.2 | 58.5 ± 1.8 | $F(1,56) = 7.56$ ($p = 0.008$) | $F(1,56) = 0.05$ ($p = 0.83$) | $F(1,56) = 4.12$ ($p = 0.05$) |
| | Large | 53.2 ± 1.7 | 49.9 ± 1.3 | | | |
| Pupil area | Small | 1.004 ± 0.005 | 0.956 ± 0.005 | $F(1,250) = 32.85$ ($p < 10^{-4}$) | $F(1,250) = 96.4$ ($p < 10^{-4}$) | $F(1,250) = 0.11$ ($p = 0.74$) |
| | Large | 1.038 ± 0.006 | 0.984 ± 0.005 | | | |

| | | | | | | |
|---|---|---|---|---|---|---|
| V4 spike rate | Small | 0.851 ± 0.007 | 0.825 ± 0.006 | F(1,22516) = 16.08 ($p < 10^{-4}$) | F(1,22516) = 21.34 ($p < 10^{-4}$) | F(1,22516) = 0.5 ($p = 0.48$) |
| | Large | 0.892 ± 0.007 | 0.886 ± 0.006 | | | |

**Supplementary Table S5: Single-trial decay/rise constant (τ) for behavior (sensitivity, d'), physiology and V4 neurophysiology in response to reward changes.**

| | Behavior ($\tau_{d'}$, 95% CI) | | Pupil area ($\tau_{pupil}$, 95% CI) | | V4 spiking ($\tau_{neuron}$, 95% CI) | |
|---|---|---|---|---|---|---|
| | small→large | large→small | small→large | large→small | small→large | large→small |
| Monkey P ($N_{small}$ = 77; $N_{Large}$ = 86) | 11.4 (5.6, 17.3) | 1.1 (–0.1, 2.3) | 2.7 (2.2, 3.1) | 13.7 (12.4, 15.0) | 2.6 (1.5, 3.7) | 11.1 (8.4, 13.8) |
| Monkey S ($N_{small}$ = 128; $N_{Large}$ = 126) | 15.6 (8.5, 22.6) | 0.8 (–0.1, 1.7) | 5.1 (4.1, 6.0) | 8.7 (6.9, 10.5) | 1.4 (–0.4, 3.1) | 6.3 (3.4, 9.1) |

### Supplementary Figure

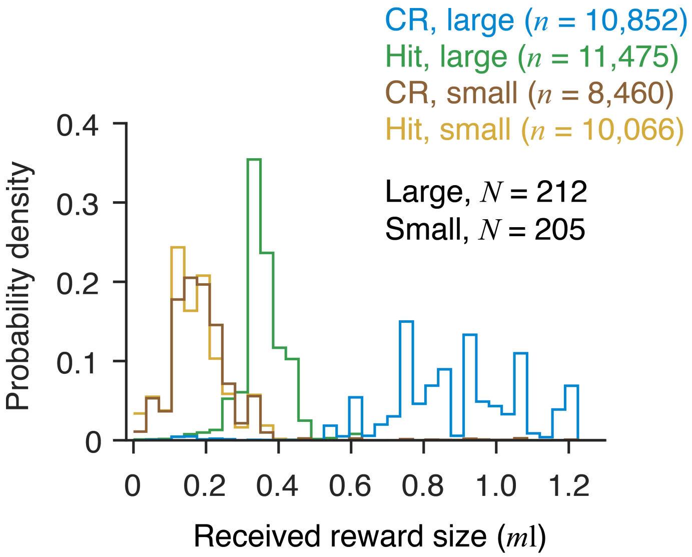


**Supplementary Fig. S1. Distribution of received rewards.** Distributions of trial-by-trial received rewards (normalized within a session) for correct responses, hits (Hit) and correct rejections (CR) across small ($N$ = 205) and large ($N$ = 212) reward blocks.

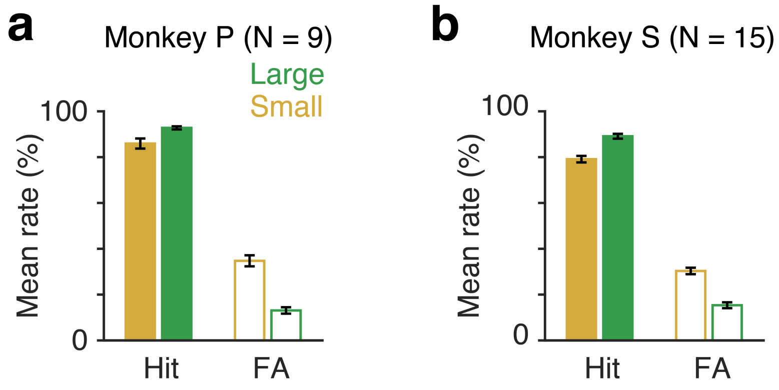


**Supplementary Fig. S2: Mean behavioral performance across sessions.** Hit and FA rates during small and large reward blocks for monkey P (N = 9 sessions). **b** Same as in **a** for monkey S (N = 15 sessions). Error bars, ± SEM.

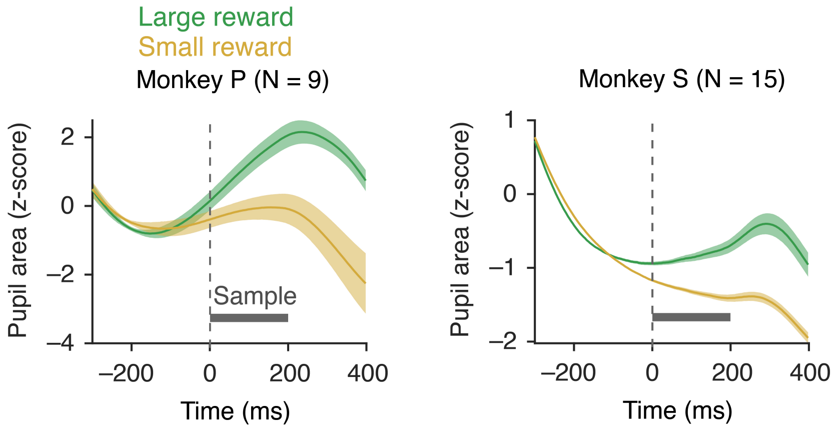


**Supplementary Fig. S3: Session averaged stimulus-evoked pupil area for small and large rewards.** Pupil areas on single trials were first aligned with the sample stimulus onset. Time course of mean pupil area within a session was z-scored with respect to the pre-sample fixation period (–400 to 0 ms) separately for two different reward size. Left, monkey P (N = 9). Right, monkey S (N = 15). Error bars, ±SEM.

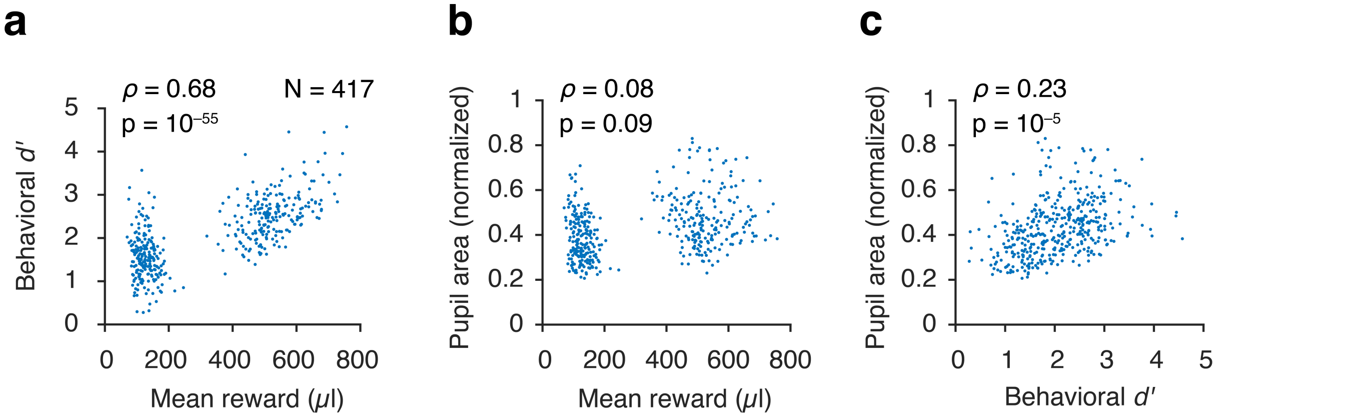


**Supplementary Fig. S4: Correlations between block-by-block trial averaged behavioral d', pupil area and reward size.** **(a-c)** Distributions of mean reward size versus behavioral d' (a); reward size versus pupil area (b) and behavioral d' versus pupil area (c) across all reward-blocks (N = 417).

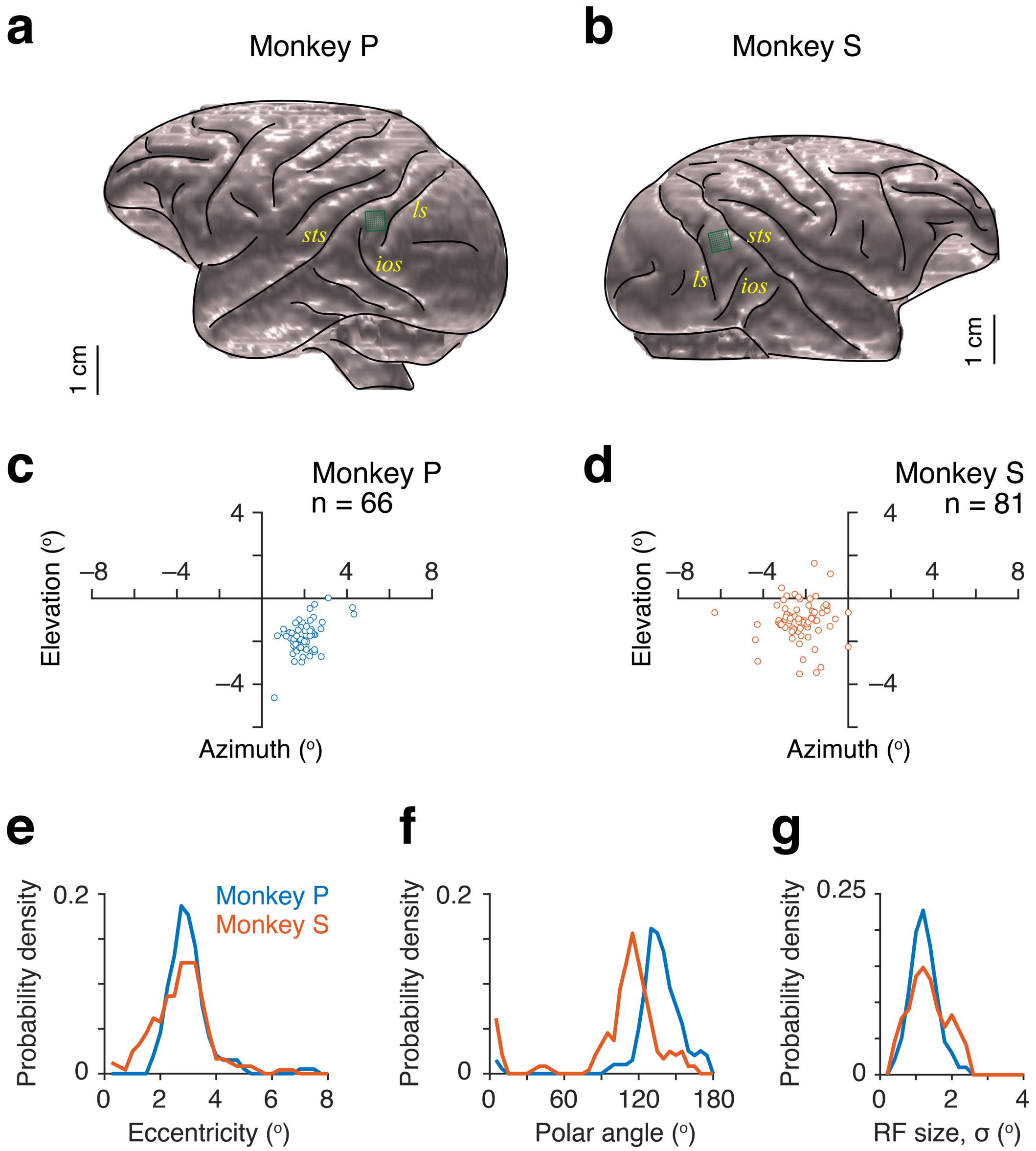


**Supplementary Fig. S5: Electrode array placements. a-b** Reconstructed brain structural MRI of monkey P (a) and monkey S (b). sts, Superior temporal sulcus; ls, lunate sulcus; ios, inferior occipital sulcus. Green box with grids, 10x10 electrode array. **c-d** Spatial receptive field (RF) centers of all unique units (only one unit from each electrode contact) from monkey P (c) and monkey S (d). e-g Distributions of RF eccentricities (e), polar angles (f) and RF size (sigma; g) for both monkeys. There is no difference between the eccentricities ($p = 0.18$, ranksum test; e) and RF sizes ($p = 0.76$, ranksum test; g) of recorded units in the two monkeys. RF Polar angles between the two animals differed significantly ($p < 10^{-11}$, ranksum test; f).

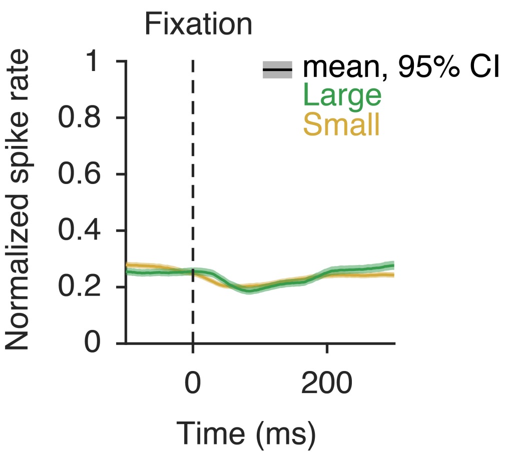


**Supplementary Fig. S6. Population peri-stimulus time histogram of spike rates of V4 units aligned to fixation.** Spike rates of each neuron (n = 970) were normalized to its peak response within 60 - 260 ms from sample stimulus onset similar to Fig. 3b-c. Error bars, 95% confidence intervals (bootstrap, $n = 10^4$). Mean spike counts over 200 ms (60 – 20 ms from fixation) did not differ across population of V4 units (mean ± sem, for all units, 5.16 ± 1.5 $s^{-1}$ (small reward), 5.51 ± 0.15 $s^{-1}$ (large reward), $p = 0.06$; for single units, 4.63 ± 0.27 $s^{-1}$ (small reward), 4.79 ± 0.28 $s^{-1}$ (large reward), $p = 0.72$; for multiunits, 5.40 ± 0.17 $s^{-1}$ (small reward), 5.82 ± 0.18 $s^{-1}$ (large reward), $p = 0.04$; ranksum test).

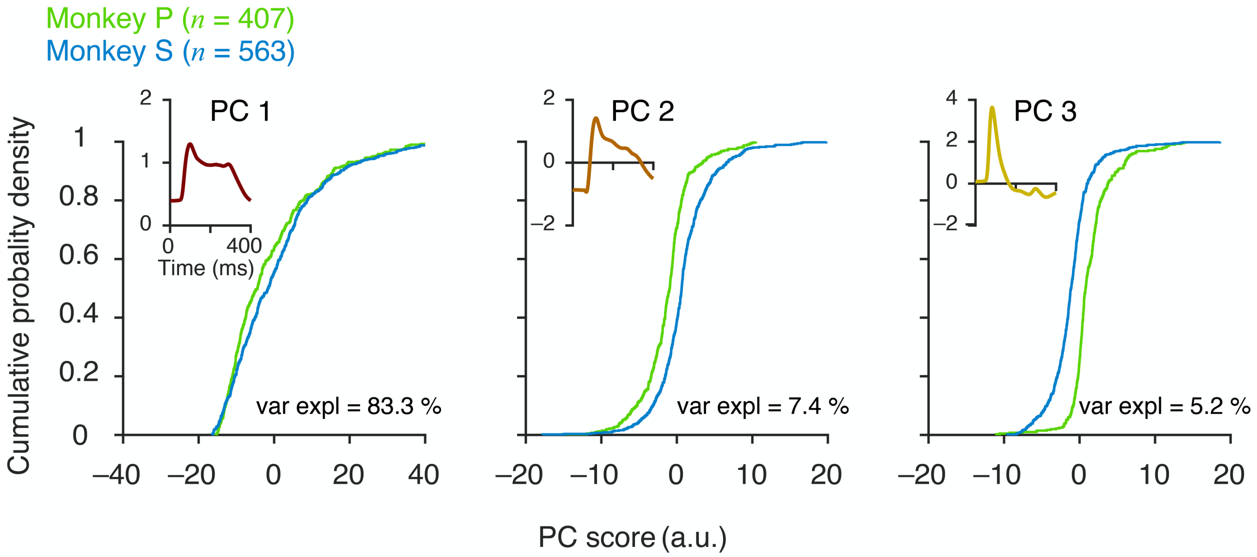


**Supplementary Fig. S7. Comparison of principal components of spike count PSTHs between animals**. Principal components analysis on spike peri-stimulus time histograms of all recorded neurons' (PSTH; 0-400 ms from sample stimulus onset during all block of trials; $n$ = 970). *Insets*, First three PCs. Bottom, Cumulative probability densities of PC scores (PC1, PC2 and PC3) for neurons in monkey P (green, $n$ = 407) and monkey S (blue, $n$ = 563). PC1 and PC2 primarily capture sustained spike response whereas PC3 associates with transient peak response. Monkey S has higher PC scores for PC1 and PC2 compared to monkey P (PC1 score, $p = 0.006$; PC2 score, $p < 10^{-23}$; Kruskal-Wallis test). In contrast, monkey P has higher PC3 scores compared to monkey S (PC2 score, $p < 10^{52}$; Kruskal-Wallis test).

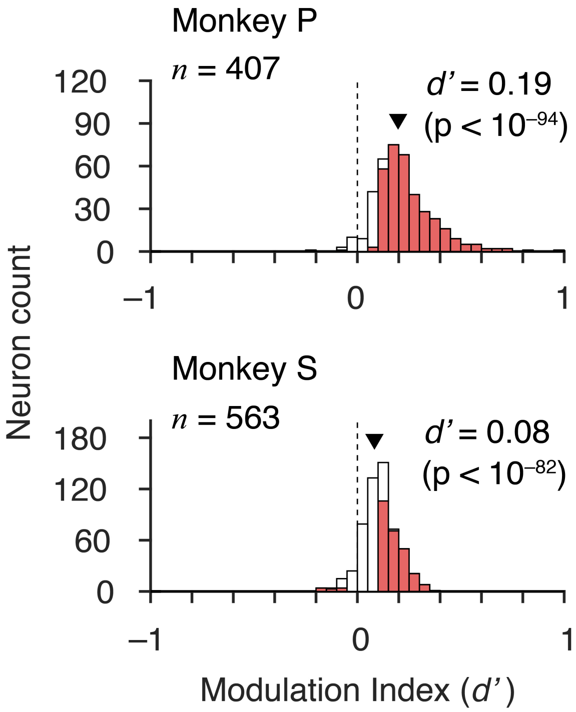


**Supplementary Fig. S8. Distribution of neuronal modulation indices (*d'*).** Distribution of modulation indices (neuronal *d'*) for monkey P (top) and monkey S (bottom). Red bars, neurons with *d'* values significantly different from zero (Monkey P, $n_{MI > 0} = 338/ 407$, $n_{MI < 0} = 2/ 407$; Monkey S, $n_{MI > 0} = 257/ 563$, $n_{MI < 0} = 11/ 563$, $p < 0.05$). White bars, non-significant MI (*d'*). Solid triangle, population MI across all units in a monkey (Monkey P, n = 407, $p < 10^{-94}$; Monkey S, n = 563, $p < 10^{-82}$, t-test).

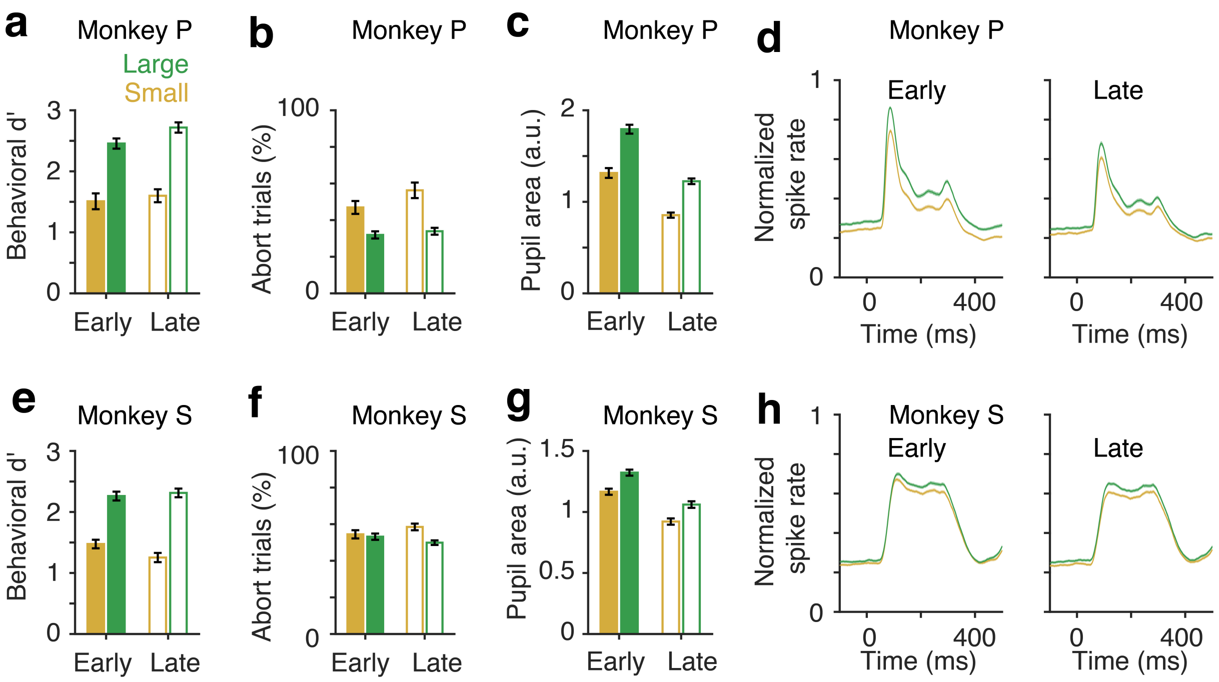


**Supplementary Fig. S9: Comparing behavioral (d'), physiological (pupil area) and neuronal modulations between early and late halves of trials within a session.** Trials were divided into two halves, early trials and late trials in every session for each monkey. **a, d** Behavioral sensitivity (d') across blocks during early and late trials with reward changes (#sessions, N = 9 for monkey P; N = 15 for monkey S). Except the reward size (small versus large), there was no significant effect of within session trial time (early versus late) on behavioral d' (monkey P, reward size, $p < 10^{-4}$, $F_{(1, 32)} = 100.94$; trial time, $p = 0.09$, $F_{(1, 32)} = 2.96$; reward-by-trial time interaction, $p = 0.4$, $F_{(1, 32)} = 0.7$; two way ANOVA). **b, g** Mean pupil area during the sample stimulus period ((#blocks, $N_{small} = 77$, $N_{large} = 86$, monkey P; $N_{small} = 128$, $N_{large} = 126$, monkey S). **c, d, h , i** PSTHs of V4 neuronal spike rates (#units, n = 407, monkey P; n = 563, monkey S). **e**, **j**, Mean V4 spike counts during sample stimuli (60-260 ms). Error bars, ± SEM.

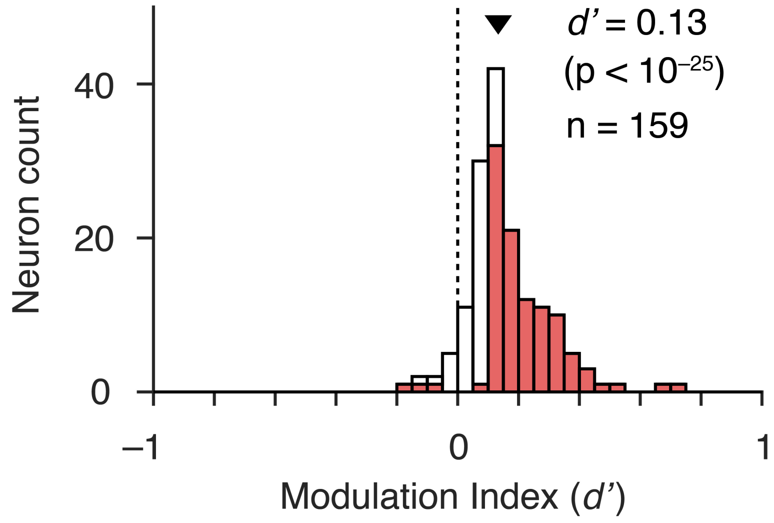


**Supplementary Fig. S10. Distribution of neuronal modulation of unique units across recording sessions.** A unique unit was randomly selected from every electrode out of a multielectrode array (96 channel) across session. Distribution of neuronal modulation (*d'*) of these unique units (n = 159) are plotted. Red bars, neurons with MI values significantly different from zero ($n_{MI>0}$ = 99/ 159, $n_{MI<0}$ = 3/ 159, $p < 0.05$). White bars, non-significant MI. Solid triangle, mean population MI across all unique units (n = 159, $p < 10^{-25}$; t-test).

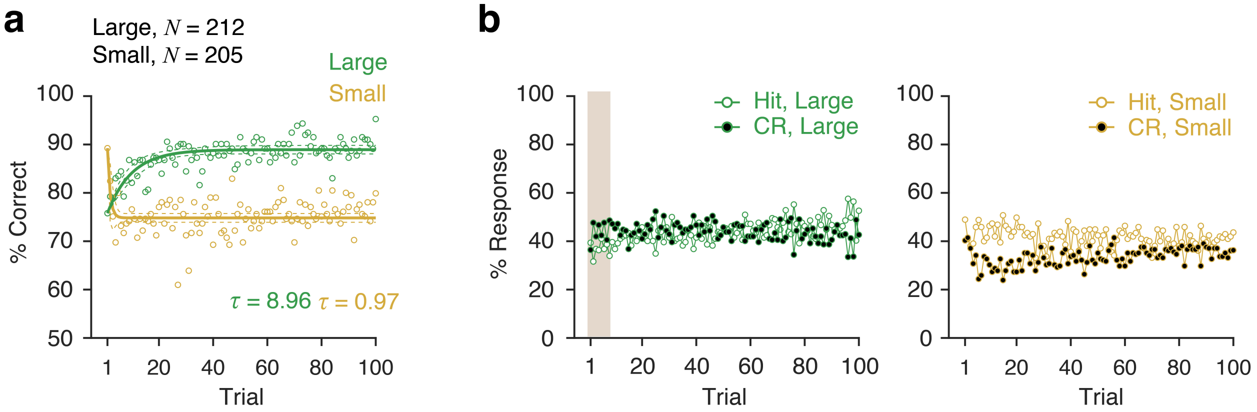


**Supplementary Fig. S11. Trial-by trial behavioral performances, percent corrects.** Block averaged trial-by-trial percentage of correct responses (**a**), hits (*left*, b) and CRs (*right*, b) for large (*N* = 212) and small (*N* = 205) reward conditions. Color bar in (**b**) highlights 1st 10 trials after the reward switches from small to large. The slow rise of percent corrects on transition from small to large reward was associated with a slow increase in hits and a transient increase in CR (b, Supplementary Fig. S12a) which led to a small increase in criterion and a slower rise of d' (Supplementary Fig. S12b and S12c). The proportion of CRs within the first 10 trials was slightly higher compared to hits for small-to-large reward switch (CR = 45.5%, hit = 37.1%, $p < 10^{-3}$, t-test). Thus, a faster rise of rewards (Fig. 5a) relative to percent correct and *d'* is primarily determined by reward size for CRs which is >2.5 fold larger than rewards size for hits (Supplementary Fig. 1).

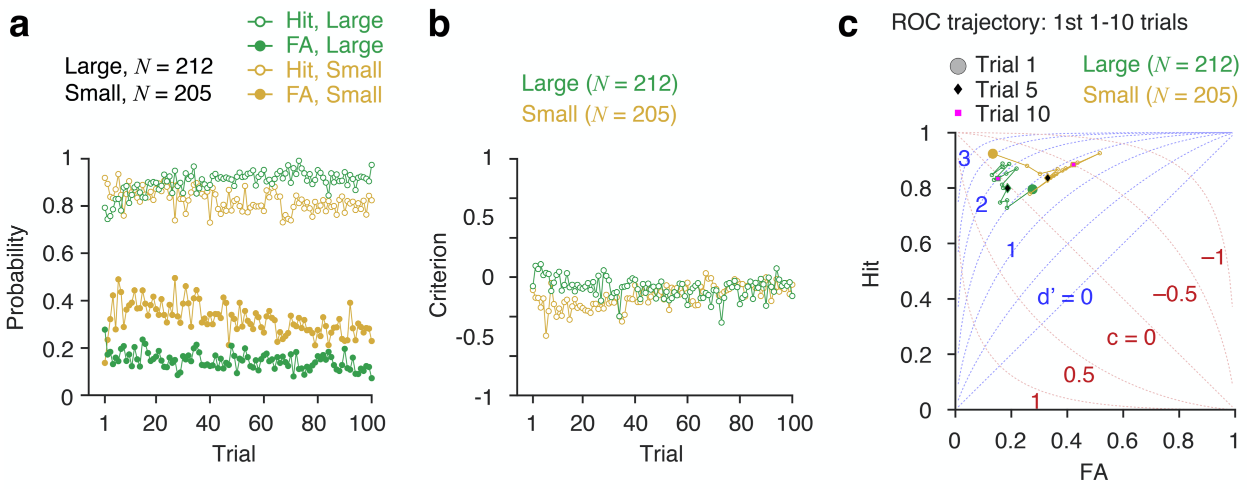


**Supplementary Fig. S12. Trial-by trial behavioral performances, response probabilities and criterion. a** Block averaged trial-by-trial probabilities of hit, miss, correct rejection and false alarm. **b** Block averaged trial-by-trial criterion for large (*N* = 212) and small (*N* = 205) reward conditions. **c** Trajectories of receiver operating characteristic (ROC) at the block transition. Filled circle, start of a block (trial 1). Diamond, fifth trial after the block transition. Square, tenth trial after the block transition.

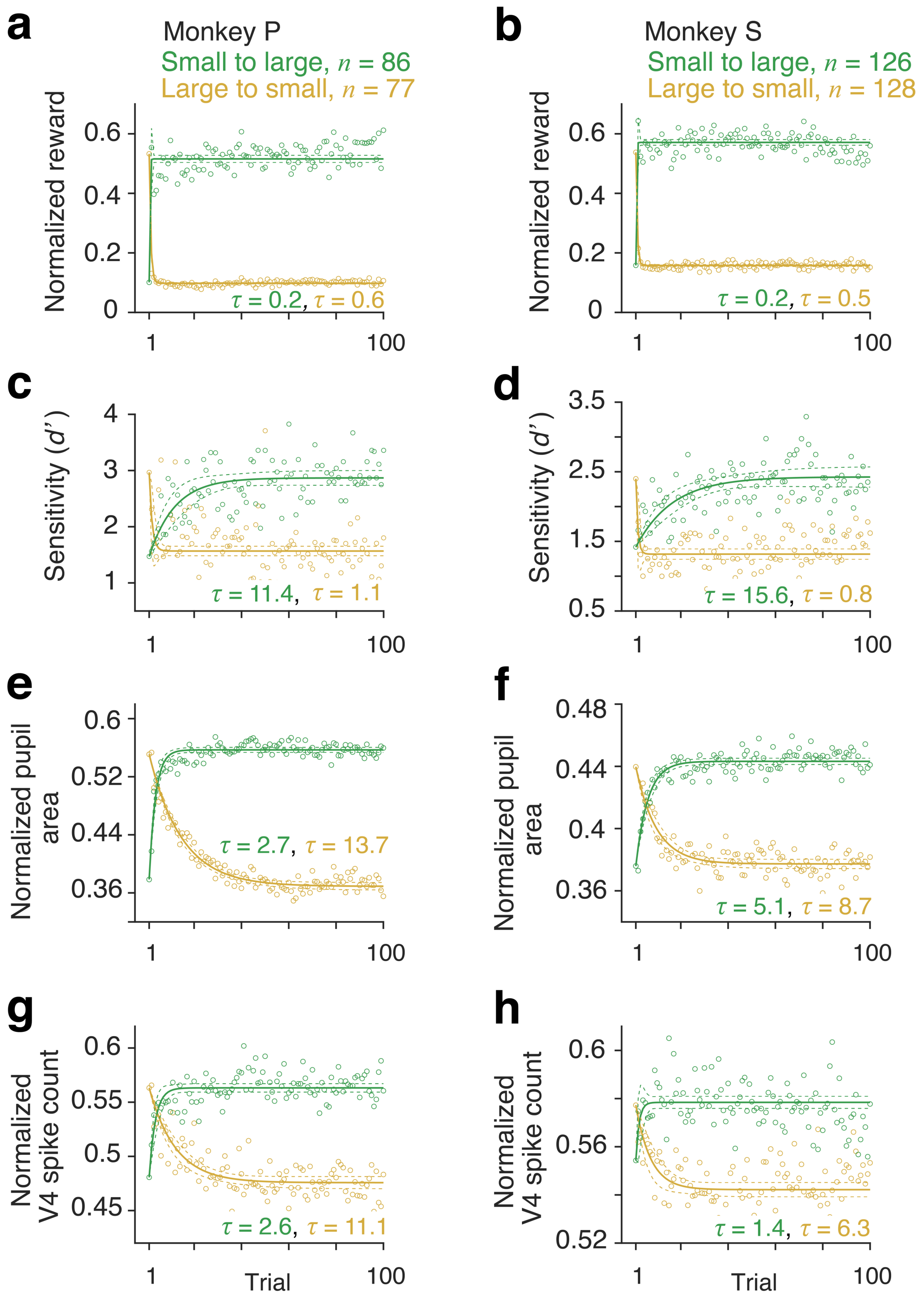


**Supplementary figure S13:** Block averaged trial-by-trial dynamics of behavioral sensitivity (d'), pupil area and V4 neuronal spiking with reward changes for monkey P (**a**, **c**, **e** and **g**; #blocks, $N_{small}$ = 77, $N_{large}$ = 86) and monkey S (**b**, **d**, **f** and **h**; #blocks, $N_{small}$ = 128, $N_{large}$ = 126). Circles, observed data. Lines, single exponential fits. τ, decay or rise constants. Trials are aligned with respect to the first correct trial following a block transition. Dashed lines, 95% confidence intervals. **a, b** Received rewards. **c, d** Behavioral sensitivity (d'). **e, f** Normalized mean pupil area during sample stimulus period. **g, h** Normalized V4 spike counts across all recorded neurons (monkey P, n = 407; monkey S, n = 563).

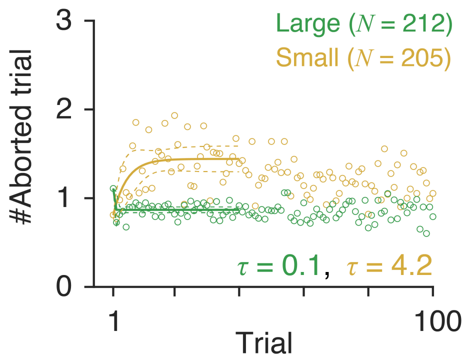


**Supplementary figure S14: Trial-by trial aborted trials.** Block averaged trial-by-trial aborted trials (fixation breaks) for large (N = 212) and small (N = 205) reward conditions. Circles, observed data. Lines, single exponential fits. τ, decay or rise constants. Trials are aligned with respect to the first correct trial following block transition. Dashed lines, 95% confidence intervals.

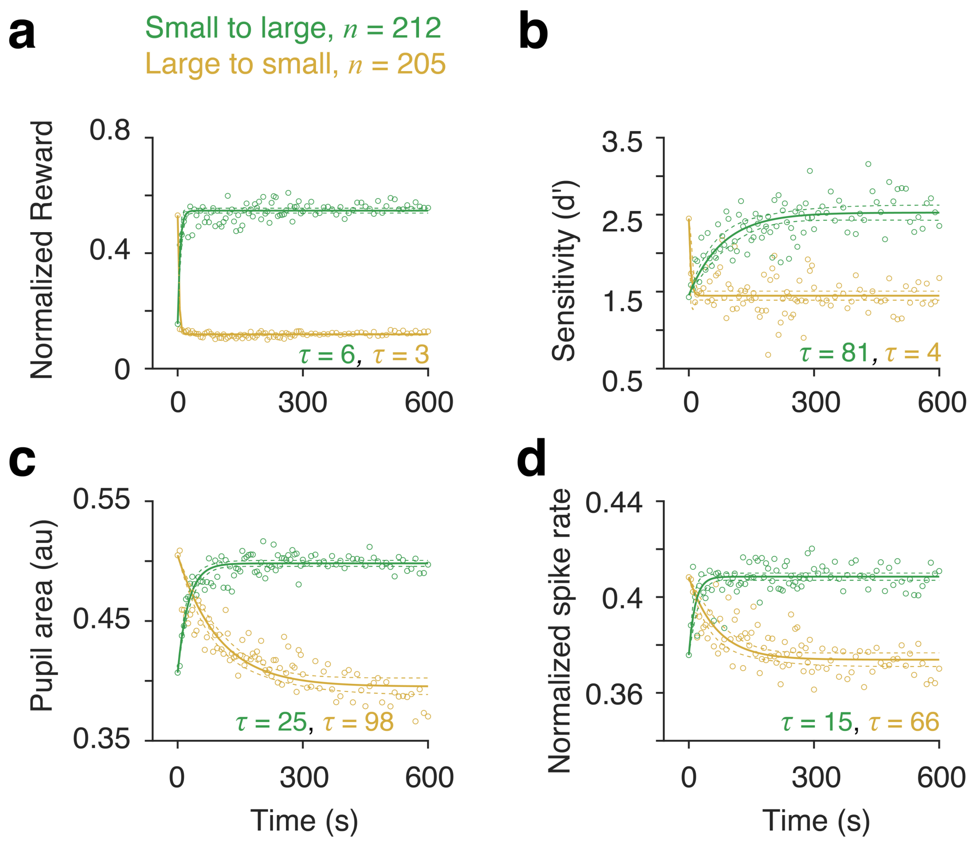


**Supplementary figure S15:** Block averaged temporal dynamics of behavioral sensitivity (d'), pupil area and V4 neuronal spiking with reward changes as a function of time (#blocks, small, 205; large, 212; two monkeys). **a** Rewards received. **b** Behavioral sensitivity (d'). **c** Mean pupil area during sample stimulus. **d** Mean normalized spike counts across blocks and neurons (n=970). τ, decay or rise constants in seconds. Trials are aligned with respect to the first correct trial following block transition. Dashed lines, 95% confidence intervals.

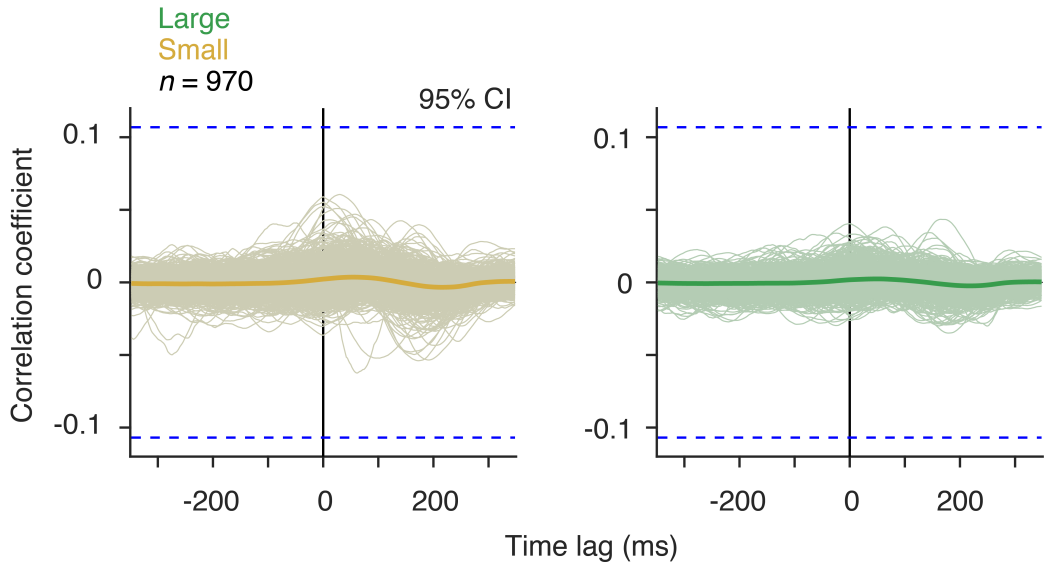


**Supplementary Fig. S16. Cross-correlation between V4 spike rate and pupil area**. *Left*, Cross-correlations (subtracted from trial shuffled values) between pupil area and single trial V4 spike rates of individual V4 neurons (*thin lines*) and averaged across neurons (*thick lines*, $n$ = 970) for small reward trials (–350 to 350 ms from sample stimulus on). Dotted line, 95% confidence interval. A positive peak at a negative lag would have indicated that changes in V4 spiking followed changes in pupil area with same sign. Cross-correlations for large reward trials similar to the *Left*.

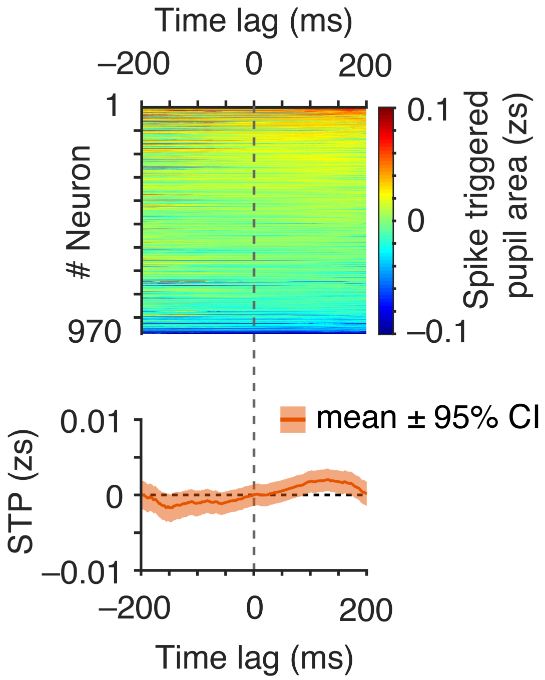


**Supplementary Fig. S17. Spike triggered (STA) averaged pupil area**. STA-pupil area (subtracted from trial shuffled values) of individual V4 neurons (*top*) and averaged across neurons (*bottom*). Spikes within 400 ms period from sample stimulus onset (0 – 400 ms) were considered. Error bars, 95% confidence intervals (bootstrap, $n = 10^4$).

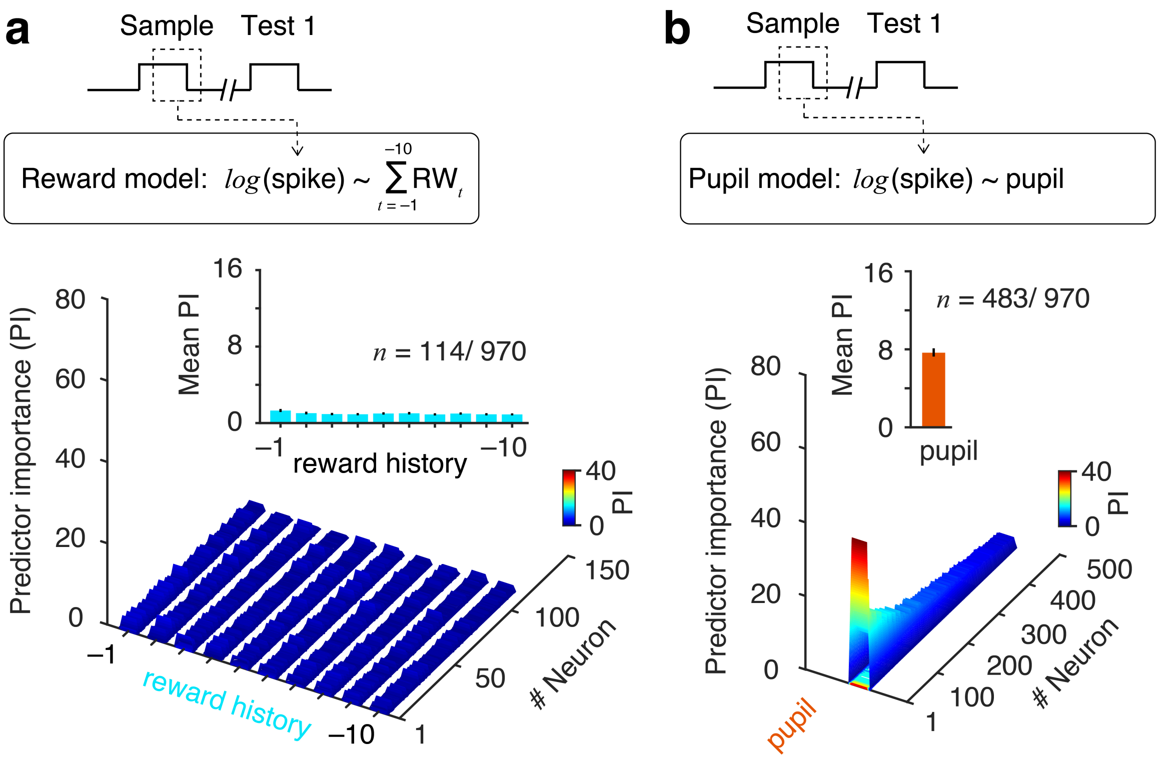


**Supplementary Fig. S18. Comparison of different reduced GLM fits in Fig. 6**. *Top* (**a, b**), schematics of the analysis time window for spike counts (60 -260 ms from sample stimulus onset) used for GLM fits. **a** Reward model, based on reward history (10 immediate past received rewards). **b** Pupil model, mean pupil area over 400 ms from stimulus onset. The same set of trials as in Fig. 6 was used. *Bottom,* (a, b), predictor importance (PIs) that measures contributions of different predictor variables estimated by absolute standardized predictor coefficient values for all the neurons that were significantly fitted (F test, $p < 0.05$). *Inset*, Mean predictor importance averaged across neurons. Error bars, 95% confidence intervals (bootstrap, $n = 10^4$).

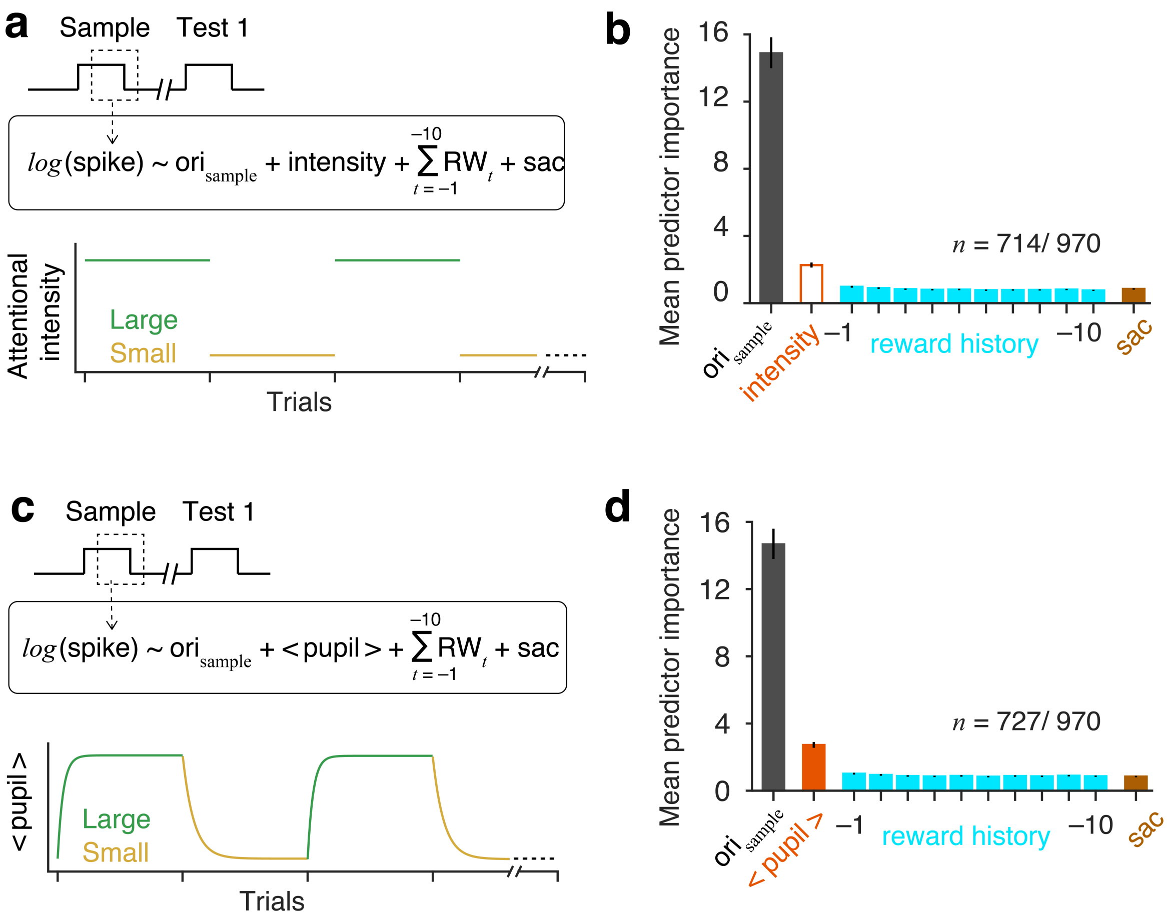


**Supplementary Fig. S19. Alternate GLM fitting of spike counts during sample stimulus. a, b** Constant attentional intensity GLM: Spike counts (60 - 260 ms from sample onset) were fit an alternate complete GLM which contained stimulus Gabor orientation of sample stimulus, constant attentional intensity (categorical variable, either ‘high’ or ‘low’; *bottom*, (a)), reward history (past 10 trials) and saccade response. Bar plot in (b) shows averaged predictor importance across neurons ($n$ = 714). **c, d** Averaged pupil area GLM: It contained stimulus Gabor orientation of sample stimulus, within session block averaged pupil area (*bottom*, (c)), reward history (past 10 trials) and saccade response. Pupil areas were averaged across large and small reward blocks separately to estimate within session single-trial-dynamics of pupil area. Thus, the variable < pupil

> has average dynamics of block transitions, but lacks single trial values. **d** Averaged predictor importance for in (c). Error bars, 95% confidence intervals (bootstrap, $n = 10^4$).

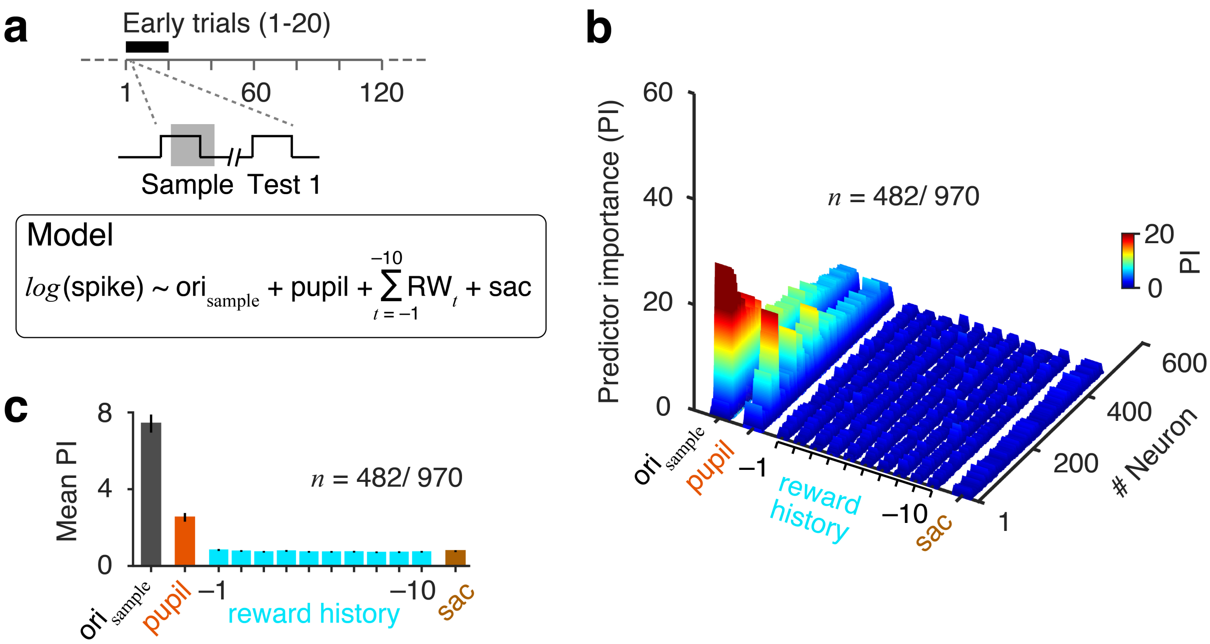


**Supplementary Fig. S20. GLM fitting of spike counts during sample stimulus for first 20 trials across all blocks. a** Complete GLM same as in Fig. 6. Spike counts are over 60-260 ms from sample stimulus onset for first 20 trials from first correct response. **b** Colormap represents predictor importance (PIs) for every neuron fitted with the complete model ($p < 0.05$, F test; $n$ = 482/970). PI measure contributions of different predictor variables estimated by absolute standardized predictor coefficient values. Neurons were sorted based on the pseudo R-squared values (Materials and Methods). **c** Averaged predictor importance across neurons ($n = 482$). Error bars, 95% confidence intervals (bootstrap, $n = 10^4$).

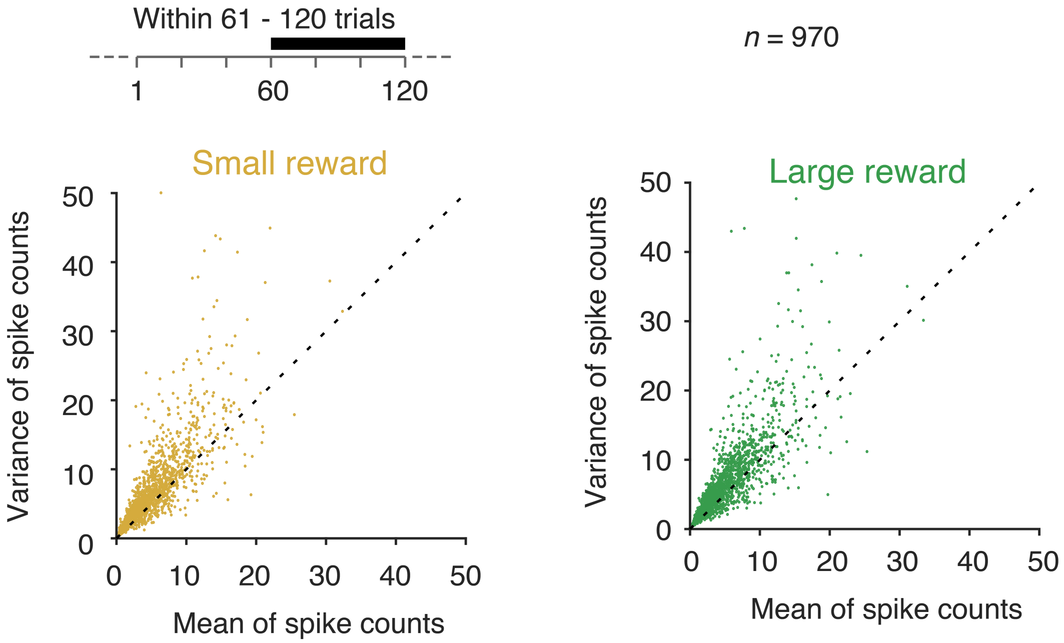


**Supplementary Fig. S21. Variances and means of spike counts**. Each marker represents the mean and variance of spike counts over a period of 200 ms (60 to 260 ms from stimulus onset) of a neuron for the same sample stimulus orientation and steady state reward/attentional intensity (trials last 60 trials in a block). Dashed line corresponds to the expected mean and variance for a Poisson count process.

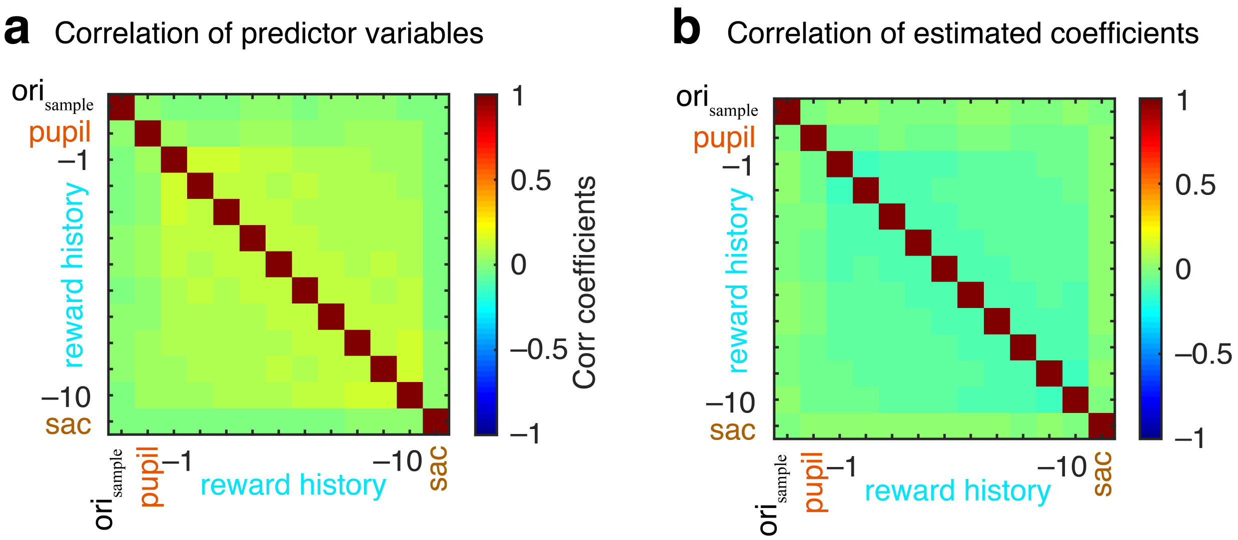

**Supplementary Fig. S22. Single trial partial correlation matrix between all predictor variable during sample stimulus period (a) and between GLM fitted predictor coefficients (b).** Dataset includes all trials in all sessions (N = 24, two monkeys, **Fig. 6**). There were no significant correlations among predictor variables or fitted coefficients ($p < 0.05$).

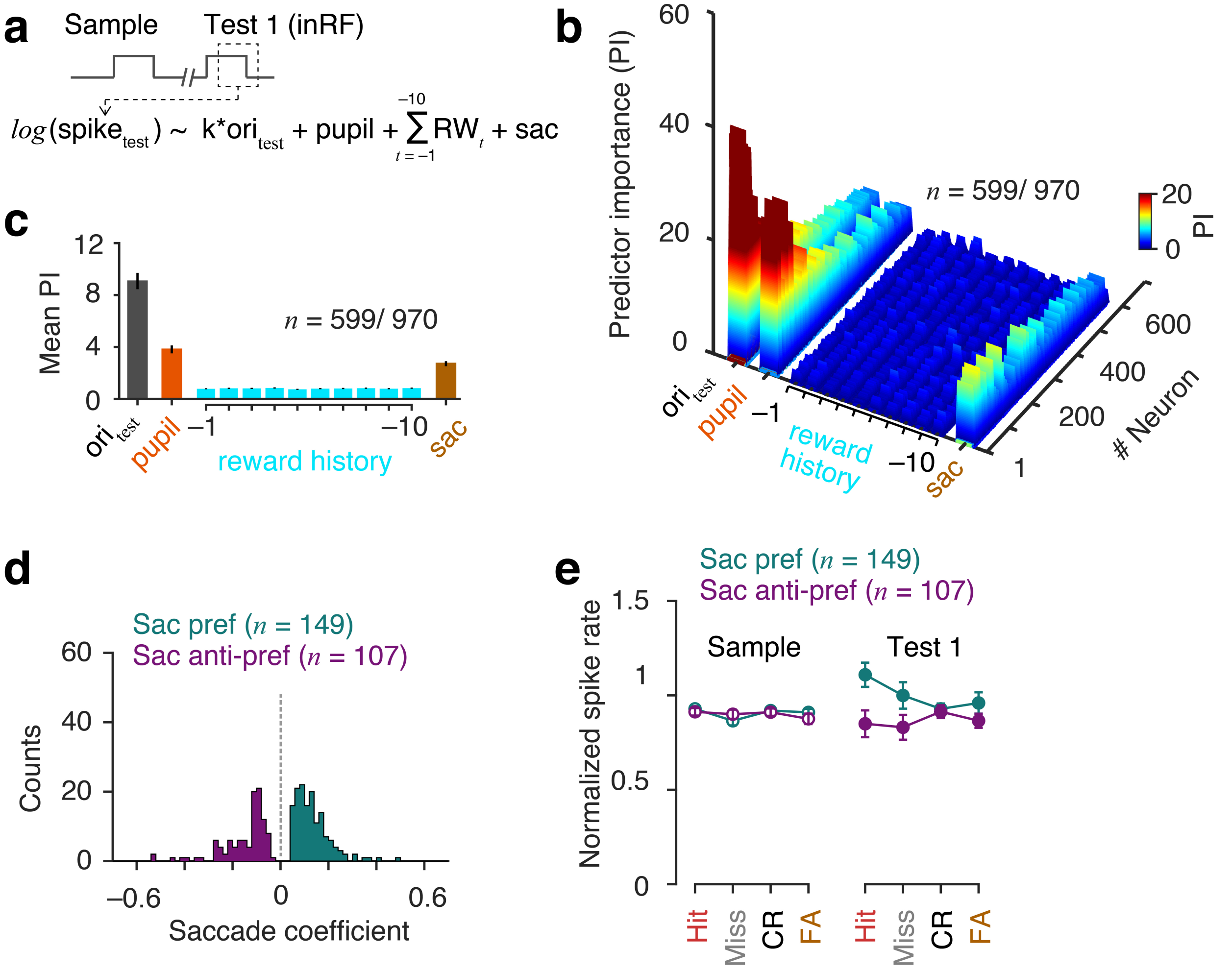


**Supplementary Fig. S23. GLM fits of spike counts during test 1 period using complete model with saccade choice as a predictor variable. a** *Top*, Saccade GLM same as **Fig. 6**. Spike counts were taken over 60-260 ms from sample onset. Predictor variables: product of test 1 stimulus orientation and neuron's orientation tuning filter, pupil area, reward history and saccade. Only a subset of trials was used for which test 1 stimulus appeared inside recorded neurons' RF and animals did not initiate any saccade before the 260 ms from test 1 on. **b, c** Predictor importance of individual neurons (b) and population average (significant fit, $p < 0.05$, F test; $n = 599/970$) (c). **d** Distribution of model fitted standardized coefficients of neurons with significant sac coefficient ($p < 0.05$, $n = 256$). Neurons with positive sac coefficient are referred as sac-preferred ($n = 149$) and with negative sac coefficient are referred as sac anti-preferred ($n = 107$). **e** Observed mean normalized V4 spike rates of saccade selective neurons (sac preferred and sac anti-preferred) for different behavior choice trials during sample and test 1 stimulus presentations (60-260 ms from stimulus onset). H: hit, M: miss, R: correct rejection, F: false alarm. Error bars, 95% confidence intervals (bootstrap, $n = 10^4$).
